# Essential role for plasma membrane glutamate transporters in stimulus intensity coding in auditory neurons

**DOI:** 10.64898/2026.04.09.717470

**Authors:** Tenzin Ngodup, Laurence O. Trussell

## Abstract

Plasma membrane glutamate transporters, also known as excitatory amino acid transporters (EAATs), serve to remove glutamate from extracellular spaces following transmitter release. Generally, this process is slow enough that EAAT activity controls the levels of background glutamate rather than the magnitude and time course of synaptic responses. We show here a striking exception to this pattern in the auditory system where neurons receive ongoing high-frequency synaptic signals. The effect of blocking EAATs on synaptic function was tested in neurons of the ventral cochlear nucleus in brain slices from mouse, focusing on auditory nerve inputs to T-stellate cells, neurons used to encode the spectrum of complex sounds. Complete block of EAATs caused a gradual accumulation of glutamate leading to a large depolarization in T-stellate cells and cessation of excitability. Partial EAAT blockade maintained resting potential but severely compromised the ability of the neurons to linearly encode the frequency of presynaptic spike activity with changes in postsynaptic firing, an essential feature of T-stellate cell function in sound intensity coding. After EAAT blockade, even a few high-frequency presynaptic spikes were sufficient to accumulate glutamate and cause repetitive postsynaptic firing lasting for tens to hundreds of milliseconds. Both glial and neuronal transporters were found to contribute to this rapid uptake necessary to maintain normal synaptic transmission. Altering the number of active auditory nerve fibers revealed that glutamate did not spill over between bouton synapses made by different nerve fibers, suggesting that synaptic boutons onto T-stellate cells restrict the diffusion of glutamate, necessitating an enhanced, local uptake activity. Notably, spike coding by the giant endbulb synapses on bushy cells was little affected by EAAT blockade, indicating a cell-type specificity to rapid glutamate uptake. Thus, rapid glutamate uptake enables the intensity coding function of neurons within the auditory system.

## Introduction

Glutamate is the principal excitatory neurotransmitter of the mammalian central nervous system. Upon release from presynaptic excitatory cells, glutamate binds to postsynaptic ionotropic receptors, generating excitatory postsynaptic potentials (EPSPs), and is subsequently cleared from the synaptic cleft through passive diffusion and reuptake by excitatory amino acid transporters (EAATs) present in both glial and neuronal membranes (Danbolt, 2001). While EAATs are critical to maintain low levels of glutamate, in most cases diffusion is so fast, and transport is so slow, that the duration of fast synaptic signaling at many central glutamatergic synapses is little affected by EAAT blockade (Isaacson and Nicoll, 1993; Sarantis et al., 1993; Takahashi et al., 1995; Turecek and Trussell, 2000; Renden et al., 2005; Graydon et al., 2014). Thus, it remains unclear whether glutamate transport plays a significant role in the millisecond time scale of neural computations in the brain.

One neural pathway which features continual glutamate exocytosis and therefore might require specialized activity of transporters is in the auditory system. For example, auditory nerve (AN) fibers may fire at 10’s of Hz in quiet conditions, and up to 400 Hz when driven by sound (Pfeiffer and Kiang, 1965; Sachs and Abbas, 1974; Taberner and Liberman, 2005). AN fibers terminate in the cochlear nuclear complex, and some of these terminals are massive, making hundreds of synaptic release sites on single postsynaptic targets, potentially accentuating the need for rapid clearance. Yet, aside from studies showing that gradual glutamate accumulation following EAAT blockade may occur at some of the giant auditory synapses, there is surprisingly little evidence supporting a major role for glutamate transport in determining how these neurons encode auditory signals (Turecek and Trussell, 2000; Renden et al., 2005; Yang and Xu-Friedman, 2009).

Indeed, many auditory nerve terminals are not ‘giant’, but conventional in structure, despite experiencing the same intensive patterns of auditory activity (Cant, 1981; Smith and Rhode, 1989a; Rubio and Juiz, 2004). We examined whether transporters play a more critical role at conventional bouton auditory synapses by studying transmission and synaptic integration at AN fiber synapses onto T-stellate cells of the ventral cochlear nucleus (VCN), a major principal cell for auditory processing. T-stellate cells contribute to sound intensity coding by effectively integrating auditory nerve inputs that are recruited as sound level is elevated (Blackburn and Sachs, 1989; Smith and Rhode, 1989b). With an increase in sound intensity, the number of active AN fibers and their firing frequency increases; the synaptic and intrinsic properties of the T-stellate cell translates this change in presynaptic activity into a linear elevation in postsynaptic firing that is steady across time, thus providing a spike code for sound level and a population code for sound spectrum (Oertel et al., 2011).

Since T-stellate cells are exquisitely sensitive to ongoing levels of synaptic activity, we speculated that they may have a more acute need for transporters to normalize activity between each synaptic response. The role of EAATs in this process was therefore tested, finding that blocking glutamate transport dramatically disrupted the ability of T-stellate cells to directly encode in their firing pattern presynaptic AN activity in the physiological range. With levels of transporter block that minimally affected baseline membrane potential, repetitive AN activity immediately led to massive pooling of glutamate, even when small numbers of AN fibers were firing. Pharmacological approaches indicated that the activity of both glial and neuronal glutamate transporters was essential to fast coding of signals by T-stellate cells. Moreover, we found that the same patterns of activity that led to breakdown of signaling in T-stellate cells upon EAAT blockade had little effect at giant terminals between AN fibers and bushy cells, synapses that encode the timing rather than intensity of sound signals. Thus, both glial and neuronal glutamate transporters play essential roles in rapid coding of sensory signals that is both cell-type and modality specific.

## Results

### Impact of full block of EAATs

We first assessed the role of transporters on membrane potential and holding current in T-stellate cells. A profound blockade of EAATs was achieved by bath application of 200 μM DL-threo-beta-benzyloxyaspartate (DL-TBOA), a concentration 4–28 times the IC50 for inhibition of EAATs 1-3 (Shimamoto et al., 1998). In current-clamp recordings from T-stellate cells the effect of block was a rapid onset of spontaneous firing, often culminating in depolarization block (Fig. 1A). The shift in membrane potential induced by EAAT blockade was 25.5 ± 4.1 mV (*n* = 5; Fig. 1B), assessed by imposing a 10 Hz filter on membrane potential after equilibration with the drug. In voltage-clamp at −70 mV, 200 μM DL-TBOA induced a large inward shift in holding current (Fig.1C, D). A cocktail of non-competitive ionotropic glutamate receptor antagonists (GYKI 53655 (50 μM) to block AMPAR and MK-801 (5 μM) to block NMDAR) reduced this shift in holding current by 70.9 ± 6.6%, indicating that much of the inward current was due to these receptors. Addition of JNJ 16259685 (1 μM) and MPEP (100 μM) to block group 1 metabotropic receptors (mGluRs) did not lead to further block of the remaining inward current (Fig. 1C–E). An additional set of experiments was performed to test whether activation of the ionotropic receptors may have obscured an mGluR component. Application of DL-TBOA after application of GYKI and MK-801 generated a very small current, and subsequent application of the mGluR antagonists did not further block this small current (Fig. 1F–G). We conclude that DL-TBOA induces a steady inward current primarily through ionotropic glutamate receptors.

**Figure 1.**
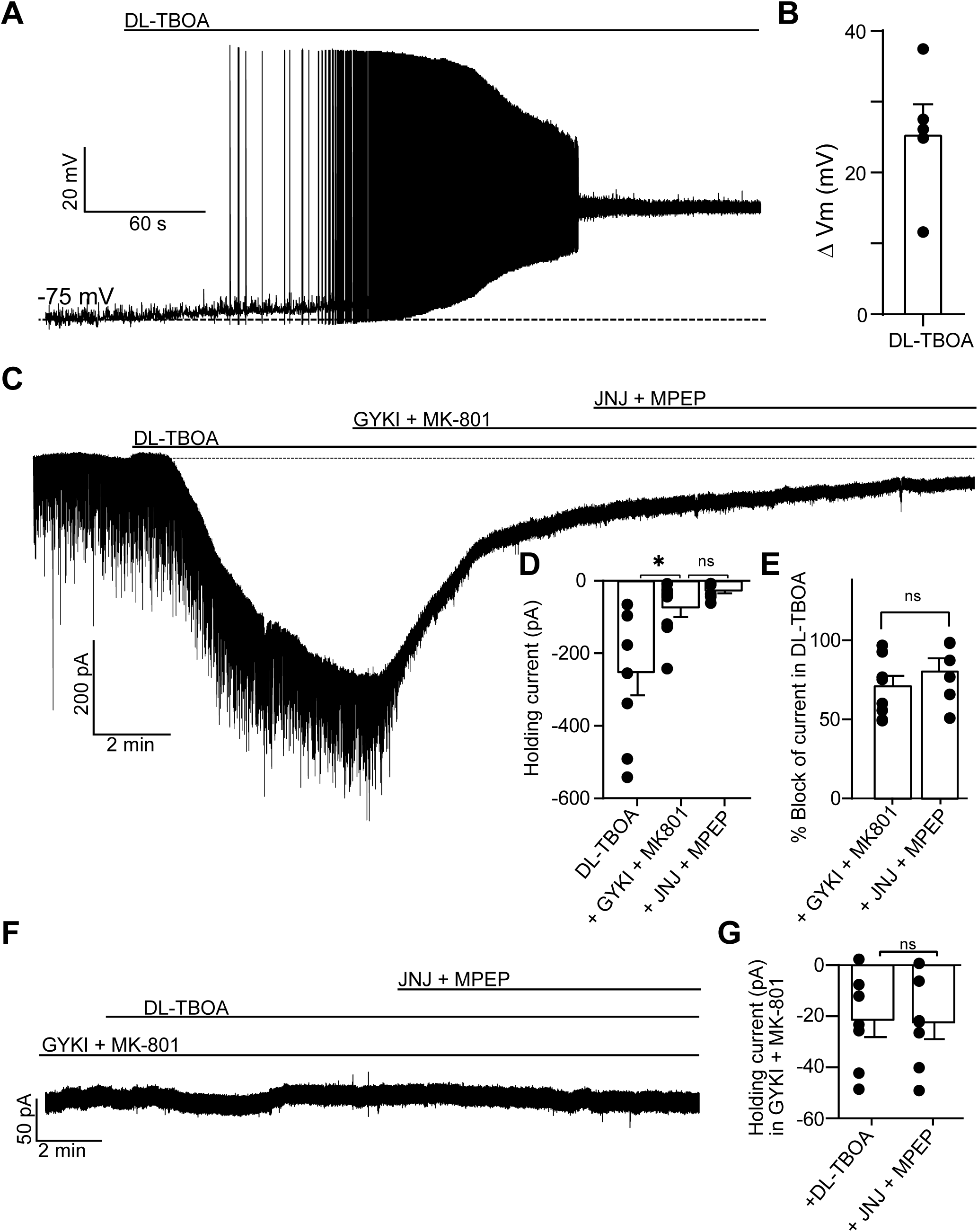
DL-TBOA induces membrane depolarization and inward current mediated by ionotropic glutamate receptors. **A**. Representative current-clamp recording from a T-stellate cell during bath application of DL-TBOA (200 μM) showing rapid onset of spontaneous firing followed by depolarization block. **B**. Summary plot of the membrane depolarization induced by DL-TBOA (25.5 ± 4.1 mV, *n* = 5). **C**. Representative voltage-clamp experiment showing a large inward shift in holding current induced by DL-TBOA. Subsequent application of ionotropic glutamate receptor antagonists (GYKI 53655, 50 μM; MK-801, 5 μM) significantly reduced this holding current. The addition of group I mGluR antagonists (JNJ and MPEP) produced no further reduction. **D**. Summary of holding current under various conditions (in pA: DL-TBOA, –250.80 ± 65.10; GYKI + MK-801, –73.25 ± 27.10; JNJ+MPEP, –26.37 ± 8.301; DL-TBOA *vs* GYKI + MK-801, *p* = 0.022, *n* = 8). **E**. Percentage block of DL-TBOA induced holding current by ionotropic glutamate receptor. AMPARs account for the majority of the inward current as mGluR inhibition provided no significant additional block (in %: GYKI + MK-801, 76.50 ± 7.47; JNJ + MPEP, 80.98 ± 7.72, *p* > 0.05, *n* = 6). **F**. Representative voltage-clamp recording in which DL-TBOA was applied in the presence of GYKI and MK-801. Prior application of receptor blockers prevented a current response upon DL-TBOA application. Subsequent application of JNJ + MPEP produced no additional change in holding current. **G**. Summary of holding current measured in the presence of GYKI + MK-801 during DL-TBOA application (–21.22 ± 6.93 pA) and after addition of JNJ + MPEP (–22.38 ± 6.16 pA, *p* >0.05, *n* = 8).

### T-stellate cells reliably encode stimulus intensity and frequency

In order to examine how transporters support coding by T-stellate cells, we characterized the sensitivity of the neurons to current or synaptic stimuli. T-stellate cells produce tonic firing in response to tones *in vivo*, a so-called ‘chopping’ firing pattern (Blackburn and Sachs, 1989; Oertel et al., 2011; Roos and May, 2012). Their firing rate increases monotonically with tone intensity, indicating that they could provide higher brain regions with information related to sound intensity (Rhode et al., 1983; Rhode and Smith, 1986; Young et al., 1988; Blackburn and Sachs, 1989; Oertel et al., 2011). A role of T-stellate cells in intensity coding is mirrored in their pattern of firing to artificial stimuli. Whole-cell patch-clamp recordings showed that T-stellate cells were able to fire with continuous spikes at a remarkably stable frequency throughout a 600-ms depolarizing current injections (Fig. 2A and B). The number of spikes during each current pulse was used to generate a frequency-*vs*-current intensity plot (Fig. 2C) and these revealed that the firing rate increased linearly with depolarizing current, only showing evidence of saturation above ∼ 500 pA. These results support previous work suggesting that T-stellate cells are intrinsically well-suited for encoding stimulus intensity (Oertel, 1983; Fujino and Oertel, 2001; Xie and Manis, 2017).

**Figure 2.**
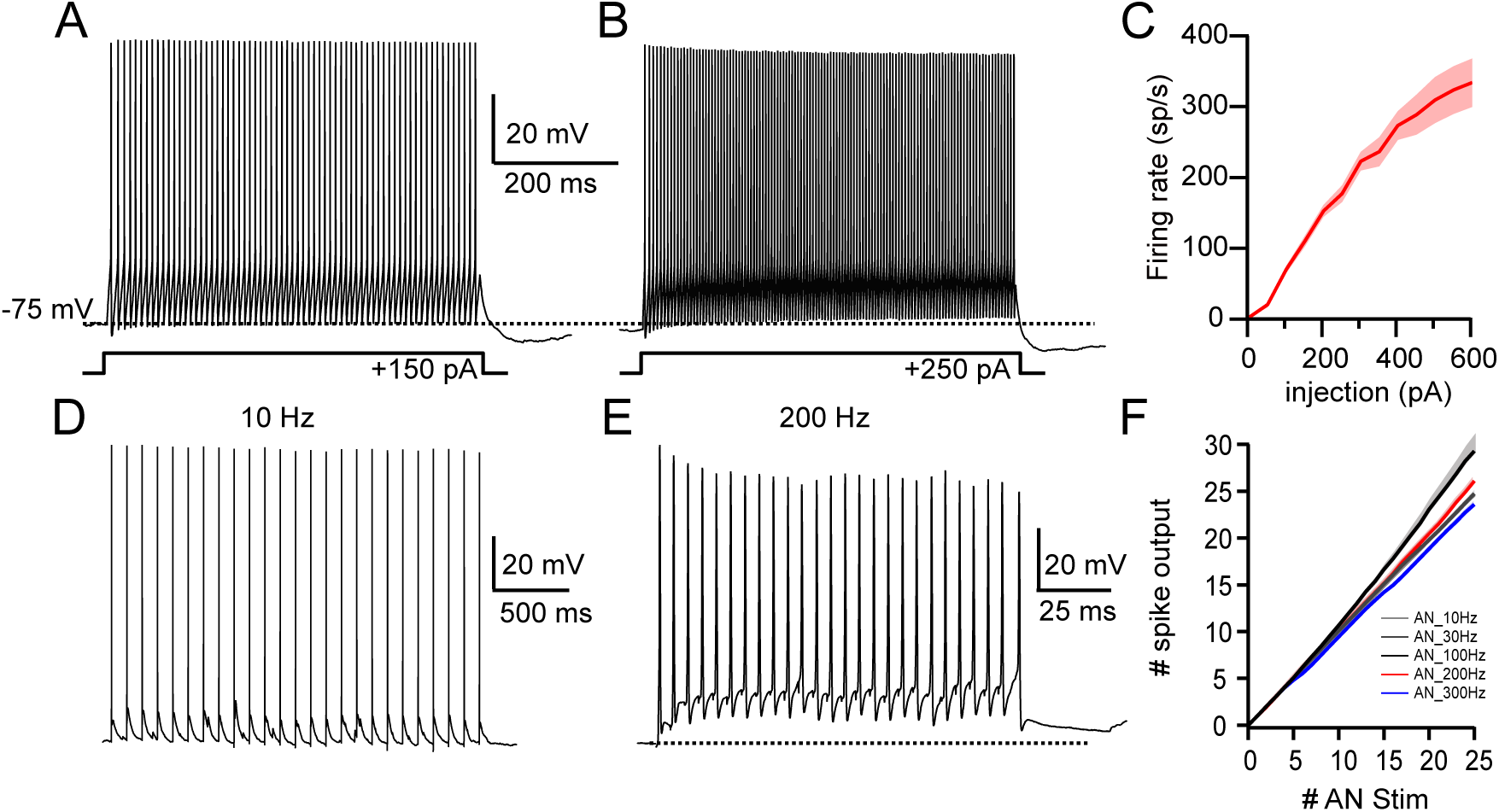
T-stellate cells reliably encode stimulus intensity and presynaptic firing rate. **A, B**. Representative current-clamp recordings from a T-stellate cell showing sustained firing during a 600-ms depolarizing current injection of +150 pA (**A**) and +250 pA (**B**). **C**. Frequency-intensity (F–I) plot showing that firing rate increases linearly with depolarizing current injection (*n* = 16). **D**, **E**. Representative current-clamp recordings from a T-stellate cell in response to a train of AN stimulation at 10 Hz (**D**) and 200 Hz (**E**). **F**. Plot showing the linear input-output relationship between presynaptic stimuli and spike output across stimulus frequencies (*n* = 6). Inhibitory glycinergic and GABA_A_ receptors were blocked using selective antagonists in all recordings.

Elevation in sound intensity increases spike rates in AN fibers. To assess the ability of T-stellate cells to encode AN spike rate, we stimulated AN fibers with a 25-pulse train at different frequencies of 10, 30, 100, 200, and 300 Hz. In all experiments, inhibitory glycinergic and GABA_A_ receptors were blocked using selective antagonists (see materials and methods). Under control conditions, each EPSP generated a single spike in T-stellate cells regardless of the stimulus frequency, resulting in a linear input-output relationship (Fig. 2D–F, *n* = 6, Fig. 3A, D). The timings of the spikes during the stimuli were used to generate raster plots of firing (Fig. 3A, D), and the number of spikes during each stimulus train was used to generate cumulative spike output plots (Fig. 3C, F). Under control conditions, T-stellate cells responded with a linear cumulative spike output to shocks of trains at different frequencies to the AN root. This result shows that the T-stellate cells fired almost exclusively during the stimulus period, rapidly terminating their spiking at the end of each stimulus train. Occasionally, we observed a few delayed spikes following higher frequency stimulations, as observed previously (Xie and Manis, 2017). This rapid termination of spiking at the end of the stimuli is vital not only for faithfully encoding sound intensity but also for preserving the precise pattern of complex sounds that characterize vocalizations (Blackburn and Sachs, 1990). Thus, rapid termination of synaptic transmission following each presynaptic signal must be exceptionally robust to render response profiles linear with the intensity and duration of presynaptic activity.

**Figure 3.**
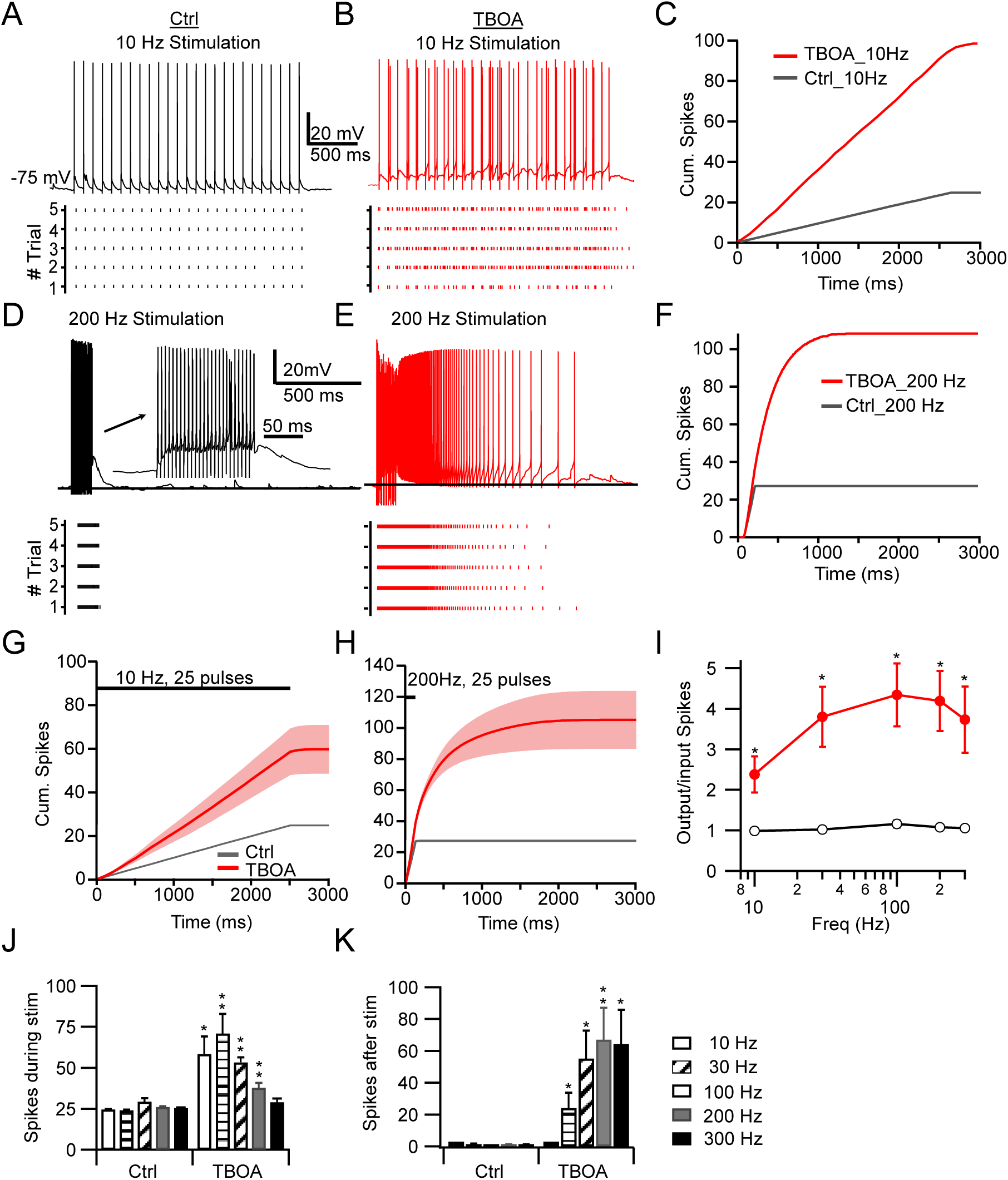
Glutamate transporters are required for precise encoding of auditory nerve activity in T-stellate cells. **A**, **B**. Representative current-clamp recordings from a T-stellate cell in response to a 25-pulse train AN stimulation at 10 Hz under control conditions (A) and in the presence of DL-TBOA (25–50 μM). Raster plots below show spike timing across trials. **C**. Cumulative spike output at 10 Hz for the same cell in A and B. Application of DL-TBOA increased spike output compared to control. **D**, **E**. Representative current-clamp recordings and raster plots of a T-stellate cell in response to a 25-pulse, 200 Hz AN stimulation under control conditions (**D**) and in DL-TBOA (**E**). DL-TBOA led to prolonged spiking following AN stimulation. **F**. Cumulative spike output at 200 Hz for the same cell in D and E. Application of DL-TBOA increased spike output compared to control. **G**, **H**. Summary of cumulative spike output plot at 10 Hz (**G**) and 200 Hz (**H**) (*n* = 6). **I**. Plot of spike output/input ratio across frequencies. Partial transporter blockade significantly increases this ratio across all tested frequencies (*p* < 0.05, *n* = 6). **J**, **K**. Summary plot showing total spikes during the stimulus train (**J**) and after the stimulus train (delayed spikes, **K**). DL-TBOA significantly increased spiking both during and after the stimulus (10 Hz, control, 0 ± 0, TBOA, 1.12 ± 0.62; 30 Hz, control, 1.33 ± 0.42, TBOA, 24.10 ± 9.46; 100 Hz, control, 0.42 ± 0.16, TBOA, 55.24 ± 17.51; 200 Hz, control, 0.89 ± 0.33, TBOA, 67.06 ± 19.99; 300 Hz, control, 0.87 ± 0.43, TBOA, 64.34 ± 21.46, *n* = 6, paired-*t*-test, \**p* < 0.05).

### Precise coding of presynaptic activity requires EAATs

Given the significant loss of membrane potential caused by a high concentration of DL-TBOA, we used submaximal concentrations (25–50 μM) with the aim of minimizing tonic glutamate accumulation while still probing the capacity of transporters to dynamically control synaptically-released glutamate during AN activity. Under these conditions, changes in membrane potential with the drug generally did not exceed 5 mV and spontaneous spiking was minimal. When the AN root was stimulated at low rates (10 and 30 Hz) in the presence of DL-TBOA, each AN stimulus resulted in at least twice as many postsynaptic spikes (Fig. 3B, C, J), an effect that could be consistent with a small reduction in spike threshold due to background transmitter. Unexpectedly, at higher stimulation rates (100-300 Hz), spiking continued long after the end of period of stimulation, lasting for hundreds of milliseconds (Fig. 3E, F, K). When we determined the ratio of the number of presynaptic spikes triggered in the AN to spike number emitted by the postsynaptic cell, we found a ratio of 1 at all frequencies under control conditions (Fig. 3I). However, after partial transporter blockade, this ratio ranged between 2.5 and 4.5, and increased with stimulus rate, such that the close relation between stimulus number and spike output at T-stellate cell synapses was lost.

T-stellate cells receive inputs from multiple AN fibers (Oertel et al., 2011), and increasing sound intensity elevates AN firing rate and firing probability. We therefore contrasted the impact of glutamate transporters when subsets of AN fibers were active *vs* when all fibers were active. First, we confirmed that we could reliably activate different numbers of AN fibers onto T-stellate cells. To do this, AN fibers were serially activated by increasing the shock duration to the AN root, and T-stellate cells were recorded in voltage-clamp to monitor the resulting excitatory postsynaptic currents (EPSCs). Shocks of various durations ranging from 10 μs to 200 μs were delivered to recruit different numbers of AN fibers onto T-stellate cells. Varying stimulus duration rather than varying stimulus voltage proved to be a more reliable means for stable, trial-to-trial activation of subsets of fibers (Lu and Trussell, 2007). When AN fibers were activated with brief shocks (10 μs), we observed small evoked EPSCs with short latencies, which increased in amplitude with longer shock durations, resulting in a graded increase in evoked EPSC amplitudes (Fig. 4), thus demonstrating electrical recruitment of additional axonal inputs to T-stellate cells. The total charge of synaptic responses as durations increased showed discrete steps (Fig. 4B), indicating that T-stellate cells receive 4 ± 2 AN fiber inputs (*n* = 7), consistent with previous studies (Ferragamo et al., 1998; Cao and Oertel, 2010; Xie and Manis, 2017).

**Figure 4.**
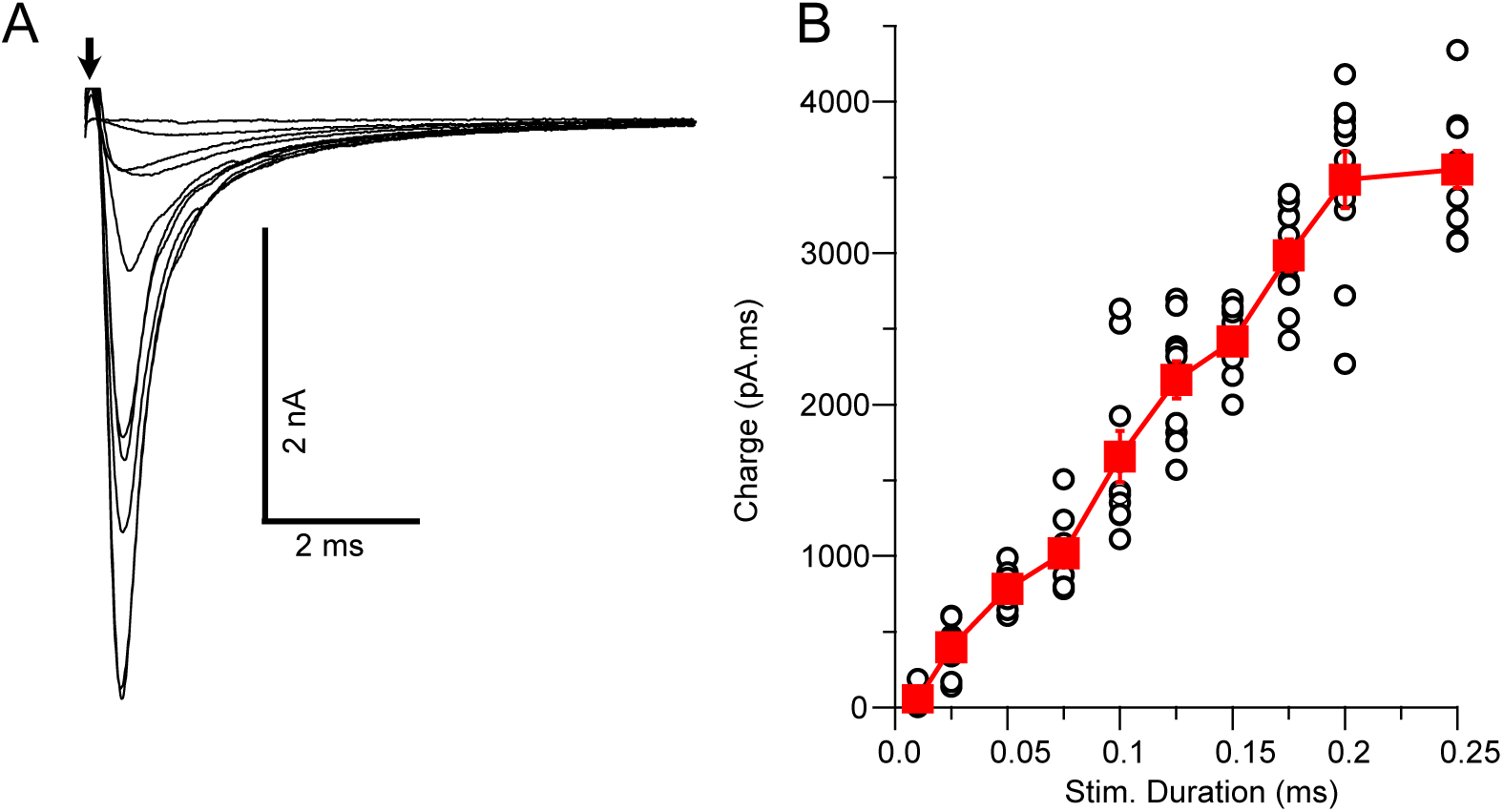
Recruitment of auditory nerve (AN) inputs to T-stellate cells. Shocks of increasing duration delivered to AN root evoked excitatory postsynaptic currents (EPSCs) that increased in a graded manner with stimulus duration. **A**. Representative voltage-clamp recording of EPSCs in a T-stellate cell evoked by AN root stimulation. EPSC amplitudes increased in a graded manner with increasing stimulus duration. **B**. Total synaptic charge plotted as a function of stimulus duration for the cell shown in A. Discrete increments in synaptic charge reflect the recruitment of additional AN fiber inputs.

Using this stimulus approach, current-clamp recordings were made from T-stellate cells, and the AN root was shocked with 25-pulse trains at 200 Hz. In control conditions, almost all EPSPs were subthreshold when AN root was stimulated with a train of brief shock pulses (50-75 μs), and as shock duration increased EPSPs become uniformly suprathreshold (Fig. 5A, C). This protocol was then repeated in the presence of 25–50 μM DL-TBOA (Fig. 5B). In DL-TBOA, even shocks of 50–75 μs generated EPSPs that were sufficient to generate spikes (Fig. 5B, C). Overall, DL-TBOA caused T-stellate cells to reach peak firing rates with shorter shock durations compared to control (Fig. 5C, D; Fig. 5C, duration at half-maximal stimulus: control, 0.094 ms; DL-TBOA, 0.071 ms; *n* = 6 cells, *p* = 0.0035, paired *t*-test). Furthermore, cells in DL-TBOA continued spiking for hundreds of milliseconds after the termination of the train stimulus even with submaximal shock durations (Fig. 5B, D & E). Thus, transporter blockers enhanced the sensitivity of T-stellate cells to even a few AN fiber inputs, saturating the capacity of the cells to register AN activity, and extending the period of postsynaptic activity far beyond the period of presynaptic activity.

**Figure 5.**
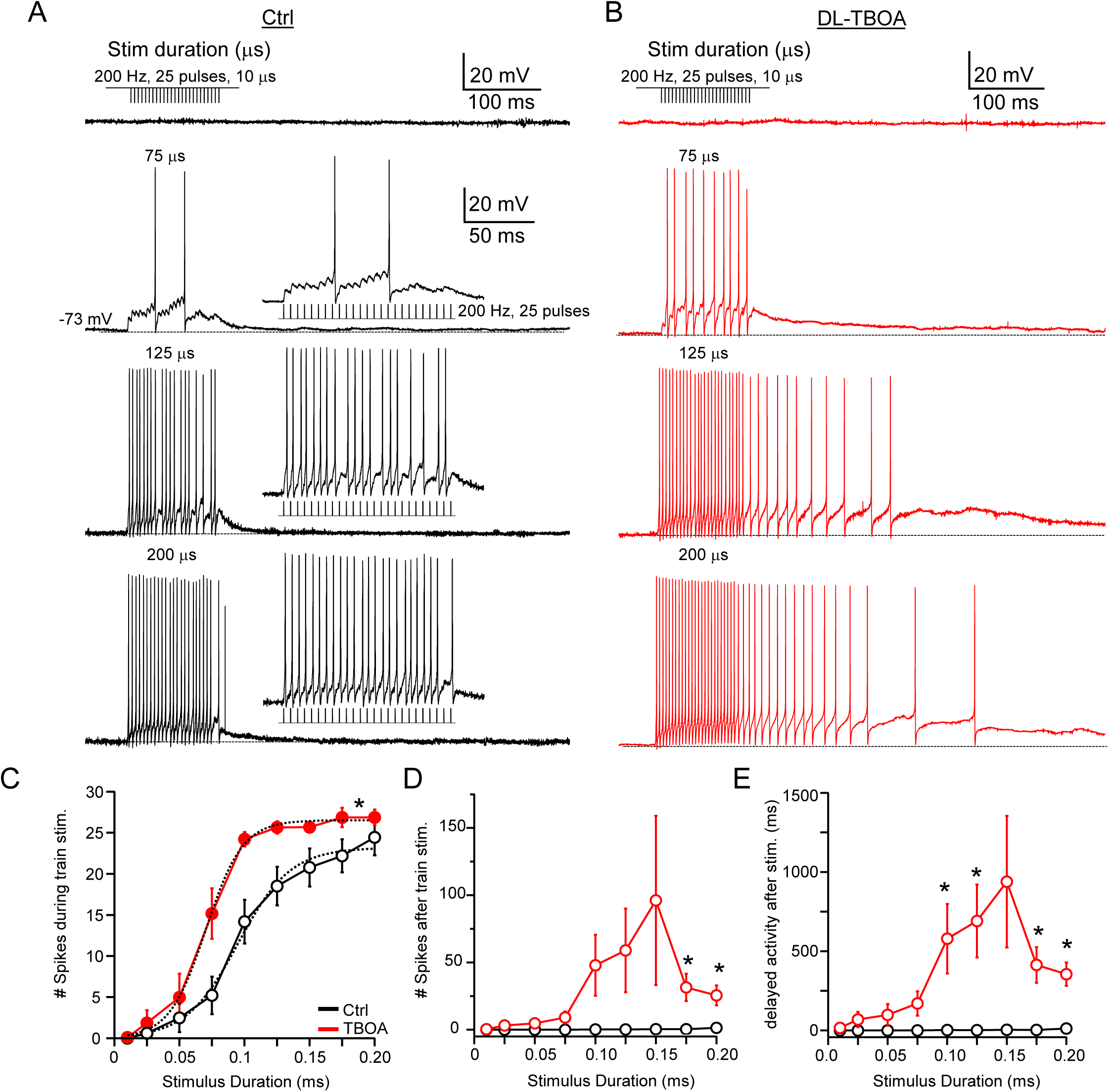
Glutamate transporter block enhances the excitability of T-stellate cells to auditory nerve input and prolongs delayed firing. **A, B**. Example current-clamp recordings from T-stellate cells during AN root stimulation (25 pulses, 200 Hz) with increasing shock duration. **A**. Under control conditions, brief shocks (50-75 μs) evoked mostly subthreshold excitatory EPSPs, while longer duration shocks (200 μs) generated suprathreshold responses and spikes largely confined to the period of the stimulus train. **B**. In the presence of DL-TBOA (25-50 μM), even brief shocks reliably evoked spikes, and longer stimuli generated more spikes during the train and for hundreds of milliseconds after the end of the stimulation. **C**. Total spikes generated during the stimulus plotted as a function of stimulus duration. DL-TBOA shifted the input-output curve to the left, reducing the stimulus duration required for half-maximal firing (ctrl, duration at half-maximal stimulus: control, 0.094 ms; DL-TBOA, 0.071 ms; *n* = 6 cells, *p* = 0.0035, paired *t*-test). **D**. Number of spikes occurring after the stimulus train (delayed spikes). Post-train spiking was minimal in control but increased significantly in DL-TBOA, especially at longer stimulus durations. **E**. Duration of delayed activity following AN stimulation. Transporter blockade produced prolonged firing that persisted for hundreds of milliseconds beyond the stimulus period. (*n* = 6, \**p* < 0.05).

### EAATs prevent rapid accumulation of glutamate

These observations made using current-clamp could reflect multiple synaptic effects resulting from the loss of transporter function. For example, at the DL-TBOA concentration used, gradual accumulation of background glutamate in the tissue could lead to a steady depolarization that enhanced hyperexcitability. More rapid accumulation during each train of synaptic activity could enhance postsynaptic currents between AN spikes, and the clearance of this accumulated glutamate could then drive additional spikes long after presynaptic activity ceased. Finally, accumulation on the submillisecond timescale might broaden individual synaptic currents leading to more rapid firing. To understand the basis of the effects of transport block on firing, T-stellate cells were voltage-clamped, and then a protocol was delivered that consisted first of an isolated AN stimulus that evoked a single EPSC (eEPSC) followed by a train of 45 stimuli at 200 Hz that generated a train of EPSCs (Fig. 6A).

**Figure 6.**
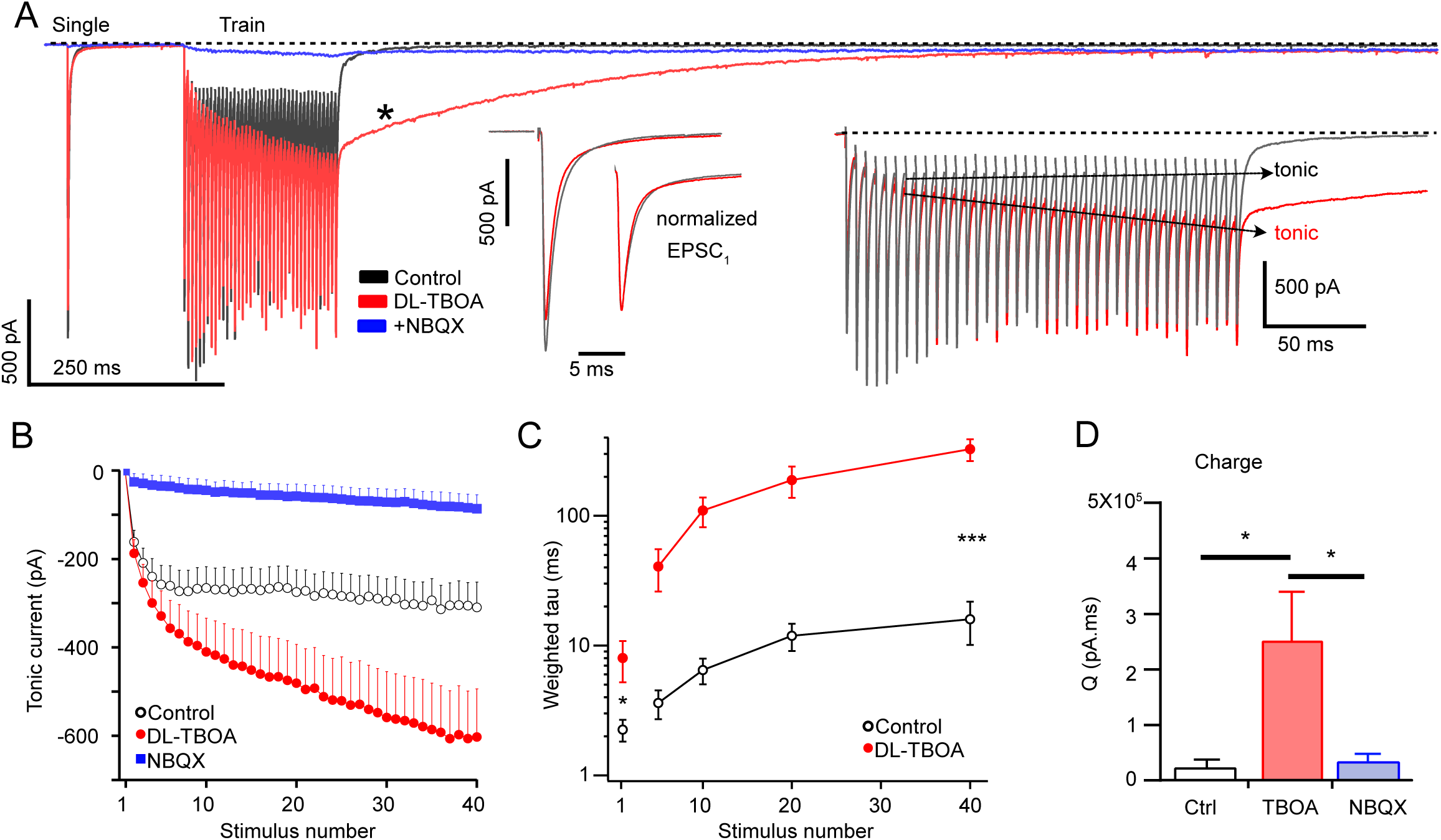
Glutamate transporters prevent glutamate accumulation during high-frequency AN activity. **A**. Example voltage-clamp recordings of excitatory postsynaptic currents (EPSCs) from a T-stellate cell evoked by a single AN shock followed by a 45-pulse train at 200 Hz. Under control conditions (black), electrical stimulation evoked a fast single EPSC (inset: middle) and a train response with minimal tonic current buildup during the stimulus (inset: right). Moreover, the train EPSC decayed to baseline within milliseconds (**C**). In the presence of DL-TBOA (red), the tonic current progressively built up throughout the stimulation (inset: right) and the decay of the EPSC train was markedly prolonged (*). These effects were blocked by AMPA receptor antagonist, NBQX (blue). **B**. Tonic current amplitude measured immediately before each EPSC, plotted as a function pulse number. Under control conditions (open circles), the tonic current reached a steady state, whereas in DL-TBOA, the tonic current progressively increased throughout the train. **C**. Summary plot showing weighted decay time constant of EPSC train as a function of stimulus number. DL-TBOA significantly slowed the decay time as the number of stimulations in a train increased. **D**. Total synaptic charge after the train across different conditions. (*p* * < 0.05, ** < 0.005, *** < 0.0005).

Under control conditions (Fig. 6A black trace), repetitive AN root stimulation caused a current buildup (‘tonic current’) between each fast eEPSC which decayed over tens of milliseconds following termination of the stimuli (Fig. 6A inset, B, C). Bath application of 25 μM DL-TBOA only slightly reduced the amplitude of single eEPSCs (Fig. 6A, left inset; mean ratio TBOA/control: 0.85 ± 0.05, *n* = 10), but caused a significant increase in tonic current during the train stimuli (Fig. 6B). We quantified this tonic current by measuring its amplitude just before each eEPSC in the train (Fig. 6A, B). While the tonic current reached a steady state in control conditions, in the presence of DL-TBOA, its amplitude began to exceed control values by the 2^nd^ or 3^rd^ stimulus in the train and more than doubled by the 40th stimulus without reaching a plateau (Fig. 6B). Strikingly, after the train, current levels decayed far more slowly in the presence of DL-TBOA, with current decay lasting for hundreds of milliseconds (Fig. 6A *). To quantify this effect, we fit the decay phase of current following the stimuli to a sum of 3–5 exponential components (see Methods). Despite the multicomponent complexity of current decline, the fastest time constant of current decay was unaffected either by DL-TBOA or by altering the number of stimuli (control, 5 stimuli, 0.55 ± 0.19 ms, 70 ± 5% of total decay, *n* = 7; 40 stimuli 0.79 ± 0.24 ms, 56 ± 5% of total decay, *n* = 6; DL-TBOA, 5 stimuli, 0.67 ± 0.17 ms, 69 ± 5% of total decay, *n* = 7; 40 stimuli, 0.91 ± 0.19 ms, 55 ± 4% of total decay, *n* = 6; F _(3, 23)_ = 0.578, *p* = 0.634, multiple comparisons test, one-way ANOVA). Nevertheless, after the fast phase of decay, later components of the EPSC declined far more slowly in DL-TBOA. Moreover, this slowing of current decay and the sensitivity to the effect of DL-TBOA was tightly linked to the number of stimuli applied during the train (Fig. 6C). For example, the weighted decay time constant for single EPSCs was increased by 3.56-fold in DL-TBOA (control, 2.25 ± 0.43 ms, *n* = 7; DL-TBOA, 8.02 ± 2.81 ms, *n* = 6; *p* = 0.048, *t*-test), while after 40 stimuli the weighted decay increased 20.5-fold (control, 15.95 ± 5.82 ms, *n* = 6; DL-TBOA, 326.50 ± 62.56 ms, *n* = 6, *p* = 0.006, *t*-test), strongly suggesting that, when transporters are blocked, increasing numbers of stimuli progressively add more glutamate to the synaptic cleft, requiring longer periods for clearance. This delayed clearance of glutamate resulted in a massive increase in charge transfer associated with synaptic stimulation, measured by integrating the current for 1 s after the last stimulus pulse in a 40-pulse train (charge (pA.ms): control, 0.14 ± 0.05 X 10^5^ *vs* DL-TBOA, 1.76 ± 0.39 X 10^5^, *n* = 11, *p* = 0.0004, paired *t*-test). Importantly, we found that both the fast EPSCs and the slow tonic current were blocked by the AMPAR antagonist NBQX (Fig. 6A, D; charge (pA.ms), DL-TBOA, 2.52 ± 0.89 X 10^5^ *vs* NBQX, 0.35 ± 0.15 X 10^5^, *n* = 4, *p* = 0.03, paired *t*-test), indicating that the slow current buildup is not due to slow-acting metabotropic signaling, but rather to repeated activation of fast-gating AMPAR by persistent transmitter. Therefore, the persistence of glutamate at AN synapses likely accounts for persistent spiking after DL-TBOA application. Passive diffusion is insufficient for complete clearance of glutamate from the synaptic cleft during high-frequency AN activity, and thus glutamate transporters are essential for controlling glutamate levels during high-frequency AN activity.

### Glutamate pooling does not result in crosstalk between different AN inputs

The ability to integrate the activity of converging AN fibers is essential to T-stellate cell function and therefore requires that different AN inputs act independently. The large degree of transmitter pooling that rapidly ensues upon EAAT blockade raises the question of whether transmitter released from one AN fiber affects the receptors underlying a different AN fiber’s terminals, that is, whether different inputs lose independence when transporters are blocked. Such a loss of independence in DL-TBOA could be manifested, for example, by a progressive slowing of the EPSC as more axonal inputs are activated. The ability to stimulate different numbers of AN fibers allowed us to test this possibility.

In voltage-clamped cells, single or train AN stimuli were delivered using either the minimal stimulus duration that still elicited an EPSC or a duration sufficient to elicit the maximal EPSC for that cell. To adequately test for such crosstalk between synapses, we chose a set of 5 cells in which the EPSC amplitude ratio upon strong *vs* weak stimulation was over 5-fold (Fig. 7E, mean, 5.40 ± 1.28-fold). This protocol was then performed in the absence or presence of DL-TBOA on each cell. Figure 7A shows overlaid responses for all these conditions. Insets illustrate overlays of the normalized responses to single stimuli to compare their durations. Compared with controls, DL-TBOA had no effect on the amplitude ratio for strong *vs* weak responses (Fig. 7E, 5.77 ± 1.55; *p* = 0.86, *t*-test).

**Figure 7.**
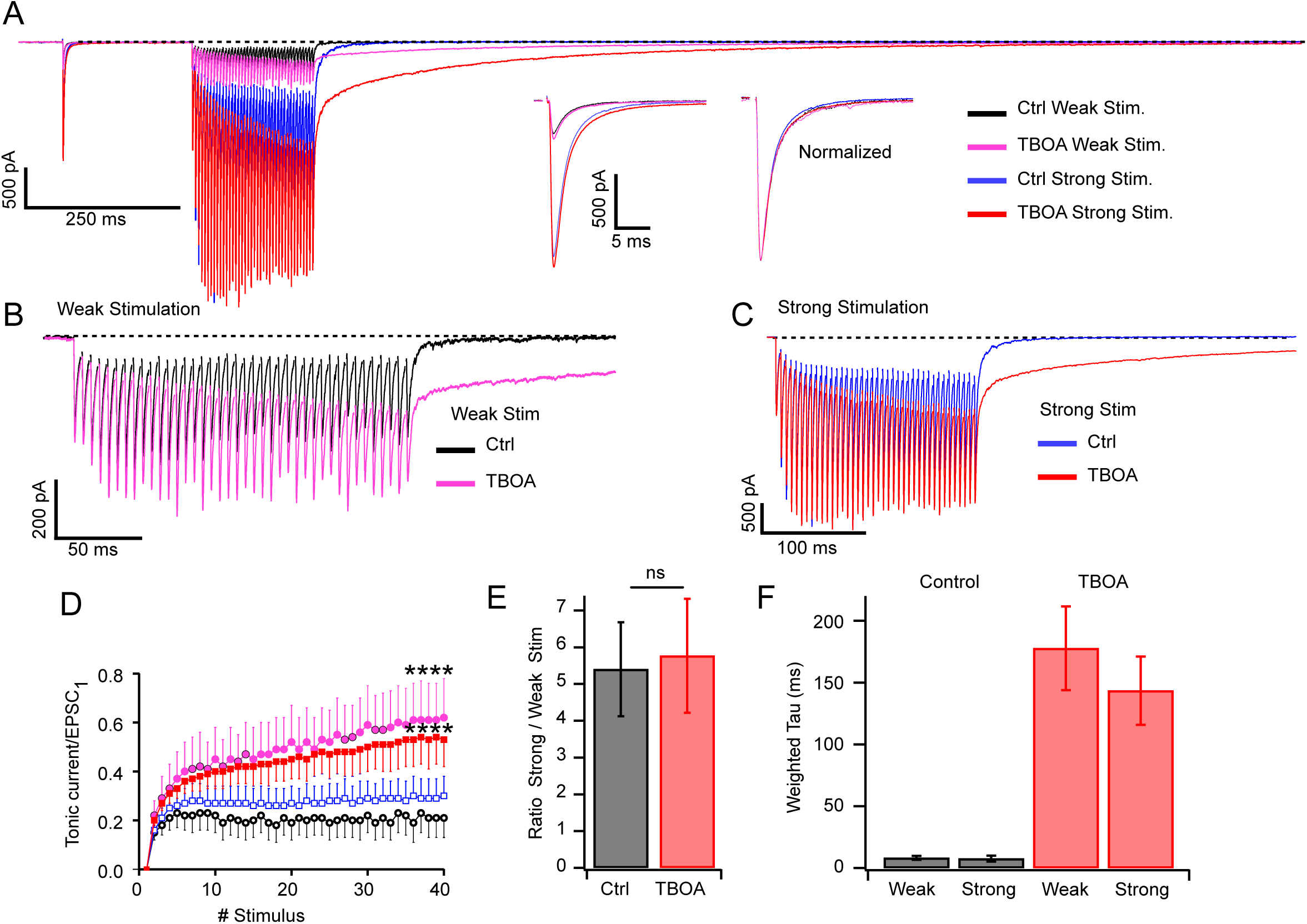
Glutamate does not spillover between auditory nerve inputs. **A**. Example voltage-clamp recordings of excitatory postsynaptic currents (EPSCs) from a T-stellate cell evoked by a single AN shock followed by a 45-pulse train at 200 Hz, delivered at weak (minimal) or strong (maximal) stimulus pulse durations before and after DL-TBOA application. Insets show single EPSC (left) and normalized EPSC (right). DL-TBOA did not alter peak amplitude of single EPSCs for a given stimulus duration. When peak-normalized all single EPSCs had similar decay times. **B**. Stimulus trains with weak (short duration) stimuli showing that enhancement of tonic current and slowed decay time after the train in DL-TBOA. **C.** As in B, but for strong (long duration) stimuli. **D.** Tonic current amplitude was normalized to the peak of the first EPSC (EPSC_1_) in a train, and plotted against pulse number during the train. While DL-TBOA enhanced tonic current, the relative enhancement was not dependent on stimulus strength. Color legend as in panel A. **E**. The ratio of EPSC_1_ for strong *vs* weak stimuli was not changed by DL-TBOA (control, 5.40 ± 1.28; DL-TBOA, 5.77 ± 1.55; *p* = 0.86, *n* = 5). **F.** Decay time (weighted tau) after stimulus trains were prolonged equally by DL-TBOA, regardless of stimulus strength.

Trains were then delivered to induce glutamate pooling. Figures 7B and 7C show for one cell a profound enhancement of tonic current and a slowing of current decay that occurred with both weak and strong stimuli. Tonic current buildup during the trains was measured just before the each EPSC and normalized to the first EPSC in each trace. While this current buildup was dramatically enhanced by DL-TBOA, when normalized to the first EPSC it was apparent that the degree of enhancement was not affected by stimulus strength (Fig. 7D, weak stimuli, control, 0.16 ± 0.04, DL-TBOA, 0.49 ± 0.13, *n* = 5, *p* = 0.01, paired *t*-test; strong stimuli, control, 0.23 ± 0.07, DL-TBOA, 0.47 ± 0.10, *n* = 5, *p* = 0.012, paired *t*-test; in DL-TBOA, weak *vs* strong stimuli, *p* = 0.44, unpaired *t*-test). Similarly, although the decay of current after the train was strongly enhanced by DL-TBOA, the decay times following the train were not different in weak *vs* strong stimulus conditions (Fig. 7F). Because a >5-fold increase in the number of active inputs, and presumably the amount of released glutamate across the whole cell, had no effect on the buildup of tonic current or on the decay time after adding DL-TBOA, we conclude that AN inputs to T-stellate cells are spatially separate from one another, or exhibit barriers to diffusion away from synaptic sites. This result stands in contrast to the progressive slowing of decay seen with increasing the number of stimuli to a given set of fibers. Thus, taken together, these observations indicate that loss of transporter function leads to pooling of glutamate but not to glutamate spread onto distant synapses and receptors.

### Both neuronal and glial glutamate transporters function during synaptic transmission

To determine whether glial or neuronal glutamate transporters contribute to glutamate reuptake at T-stellate cell synapses, we examined the effect of subtype-specific EAAT antagonists at concentrations that would saturate glial EAATs, UCPH-101 (30 μM) and DHK (200 μM), but not affect neuronal EAAT3 (Shimamoto et al., 1998; Shimamoto et al., 2004). Bath application of UCPH-101 and DHK, even at low concentrations resulted in large changes in synaptic transmission during repetitive auditory nerve stimulation. For example, Fig. 8A and Ai show an example of eEPSCs from a T-stellate cell in response to repetitive AN stimulation, in which the glial EAAT blockers markedly slowed the decay after the train (pink trace). Glial EAAT blockade increased decay time by 4.71 ± 1.27-fold and synaptic charge by 10.78 ± 3.64-fold relative to control, indicating a major role for glial transporters in glutamate clearance during sustained activity.

**Figure 8.**
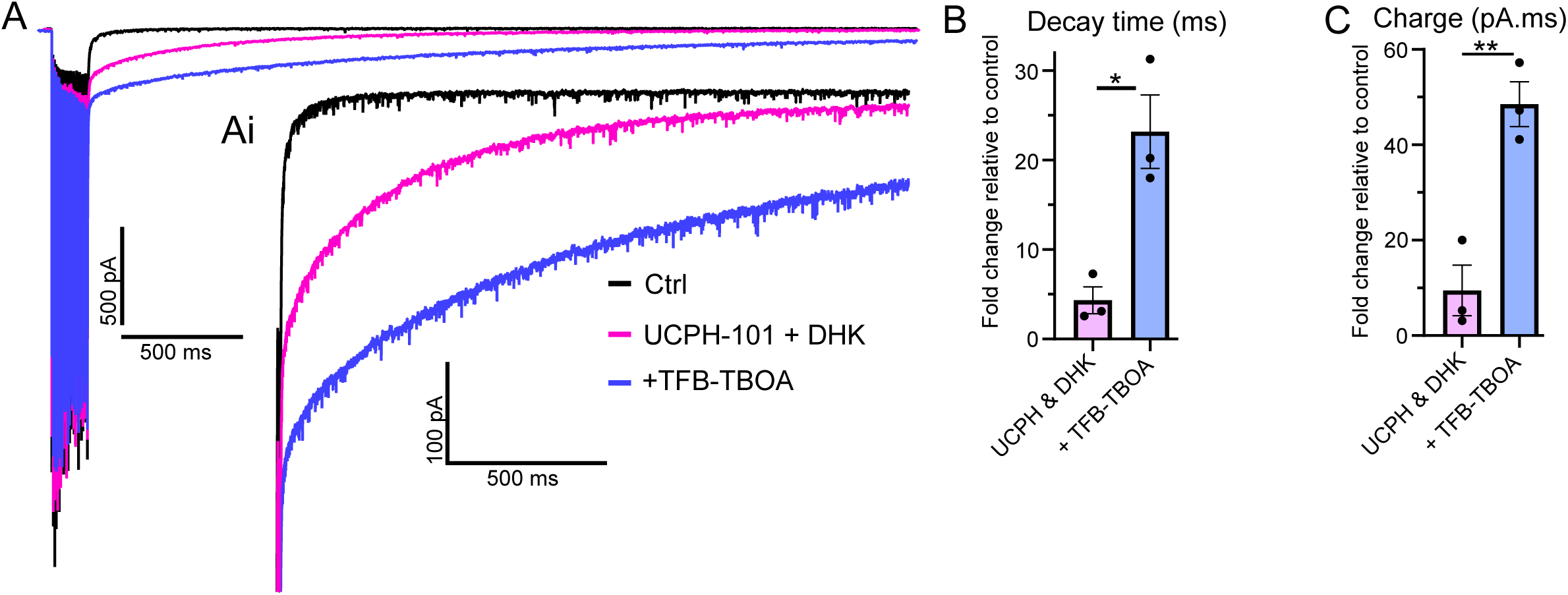
Glial and neuronal EAATs clear synaptic glutamate. **A**. Voltage-clamp recordings of response to AN stimulus train in control (black), UCPH-101 (30 μM) and DHK (200 μM) (pink), and following the addition of high-affinity blocker TFB-TBOA (2 μM). **B.** Fold increase in weighted tau after stimulus train ended, for UCPH-101 and DHK and for the subsequent addition of TFB-TBOA (UCPH-101 and DHK, 4.33 ± 1.19, TFB-TBOA, 23.17 ± 4.11, *n* = 3, *p* = 0.037). **C.** Fold change in total charge during the decay phase across these two different conditions (UCPH-101 and DHK, 9.46 ± 5.31, TFB-TBOA, 48.53 ± 4.70, *n* = 3, *p* = 0.006)

To determine whether neuronal EAATs also contribute to glutamate clearance, we performed sequential pharmacological blockade in a subset of recordings. In these cells, the non-selective EAAT antagonist TFB-TBOA (2 μM), which blocks both glial and neuronal transporters, was applied following blockage of glial EAATs with UCPH-101 and DHK. Sequential EAAT inhibition further prolonged synaptic responses during auditory nerve stimulation (Fig. 8). When normalized to baseline, combined EAAT blockade significantly increased synaptic charge and EPSC decay (charge (pA.ms), *p* = 0.019; decay time (ms), *p* = 0.029, *n* = 3, paired *t*-test, Fig. 8A, B). This effect of TFB-TBOA on synaptic transmission was significantly greater than that of UCPH-101 and DHK alone, indicating that neuronal EAATs contribute to glutamate clearance under sustained neuronal activity. Together, these findings suggest that both glial and neuronal glutamate transporters contribute to the control of glutamate clearance at the T-stellate cell synapse.

### Glutamate clearance is cell-type specific

To test whether the strong uptake dependence of AN-to-T-stellate cell transmission is cell-type specific, we recorded from bushy cells, a major principal cell in the VCN that receive giant axosomatic AN terminals, the well-known endbulbs of Held. Bushy cells were recorded in current-clamp and AN fibers were activated with trains of electrical shocks at different frequencies. At 100 Hz, bushy cells fired reliably in response to each stimulus (Fig. 9A, E). However, as documented previously, at higher frequencies (200 & 300 Hz), bushy cells failed to generate a spike to each stimulus (Fig. 9C, F) due to synaptic depression and possibly sodium channel inactivation (Yang and Xu-Friedman, 2009; Kuenzel et al., 2011; Ngodup et al., 2015), resulting in a loss of linear input-output relationship at higher frequencies (Fig. 9G). Moreover, bushy cells rapidly terminate spiking at the end of the stimulus train at all frequencies tested, such that there were never delayed spikes (Fig. 9A–D), similar to T-stellate cells. However, unlike T-stellate cells, in the presence of DL-TBOA there was virtually no effect on spike generation during and after trains of stimulation at all frequencies (Fig. 9B, D–G). Thus, endbulb synapses are able to sufficiently clear glutamate from high-intensity synaptic activity by passive diffusion without compromising postsynaptic signaling.

**Figure 9.**
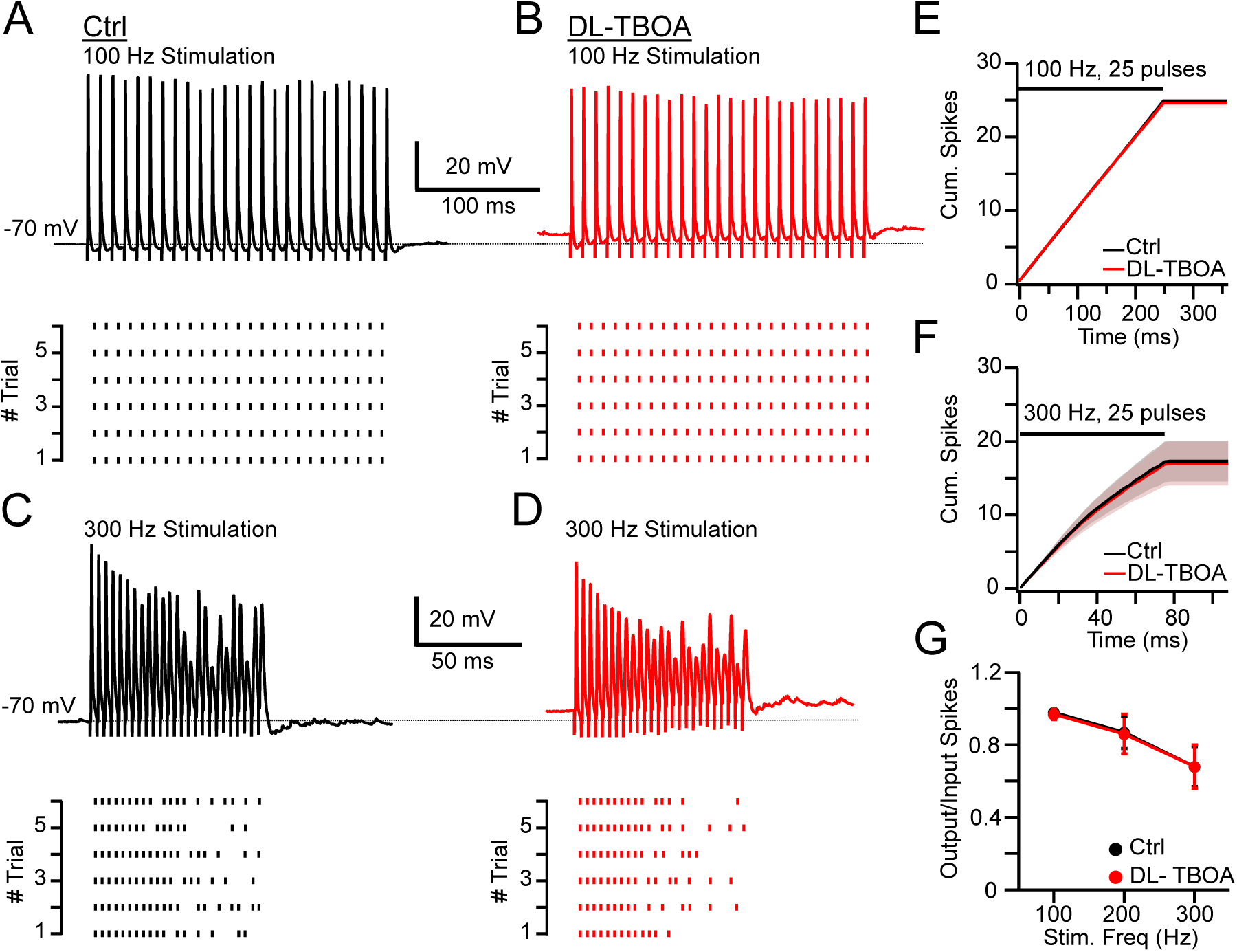
Synaptic efficacy of EPSPs in VCN bushy cells are not impacted by EAAT block. **A, B**. Current-clamp recordings of the response to 100 Hz AN stimulus trains, in control and DL-TBOA solutions, respectively. Spike raster plots beneath traces indicate consistent suprathreshold transmission. **C, D**. As in A and B, but for 300 Hz stimulus trains. EPSPs later in the trains were subthreshold but again DL-TBOA had little impact on EPSP efficacy. **E.** Cumulative spike number during 100 Hz train shows no effect of DL-TBOA. No additional spikes occurred after the train ended. **F.** As in E, but for 300-Hz stimuli. **G.** Ratio of emitted spikes in bushy cells to number of presynaptic stimuli at 100, 200, and 300 Hz showed no effect of DL-TBOA (100 Hz, control, 0.98 ± 0.01, TBOA, 0.97 ± 0.03; 200 Hz, control 0.87 ± 0.09, TBOA, 0.856 ± 0.11; 300 Hz, control 0.68 ± 0.11, TBOA, 0.68 ± 0.12, *n* = 8, *p* > 0.5).

Although no effect of transporter block in bushy cells was seen in current-clamp, we asked whether evidence of glutamate accumulation in bushy cells could be detected using voltage-clamp recordings during 200 Hz AN stimulus trains. Indeed, application of DL-TBOA resulted in a prolonged current decay and potentiation of total charge following train stimulation (Fig. 10A *). Nevertheless, when comparing the effects of the transporter blockers between bushy cells and T-stellate cells, we found that DL-TBOA had a significantly smaller effect on bushy cells than on T-stellate cells in terms of current decay and total charge transfer (Fig. 10B, charge (pA.ms); T-stellate, control, 13,616.75 ± 5,274.50 *vs* DL-TBOA, 176,319.57 ± 39,328.01, *p* = 0.0008, *n* = 11; bushy cell: control, 19,811.89 ± 13,220.82 *vs* DL-TBOA, 44,964.13 ± 23,392.42, *p* > 0.99, *n* = 8, Kruskal-Wallis test, Post hoc, Dunn’s test; weighted decay time (ms); T-stellate, control, 104.24 ± 44.58 *vs* DL-TBOA, 414.23 ± 107.17, *p* = 0.0016, *n* = 11; bushy cell: control, 81.19 ± 21.11 *vs* DL-TBOA, 173.80 ± 53.12, *p* > 0.99, *n* = 8, Kruskal-Wallis test, Post hoc, Dunn’s test). Notably the greater sensitivity of T-stellate cells *vs* bushy to transport block was in contrast to the larger amplitude of EPSCs in bushy cells in control solutions (T-stellate, 1,570.57 ± 226.37 pA, *n* = 9 *vs* BC, 4,699.55 ± 1180.01 pA, *n* = 10, *p* = 0.014, t-test). Therefore, the regulation of glutamate levels by glutamate transporters is target cell-specific, and passive diffusion plays the dominant role in glutamate clearance at the large endbulb synapse.

**Figure 10.**
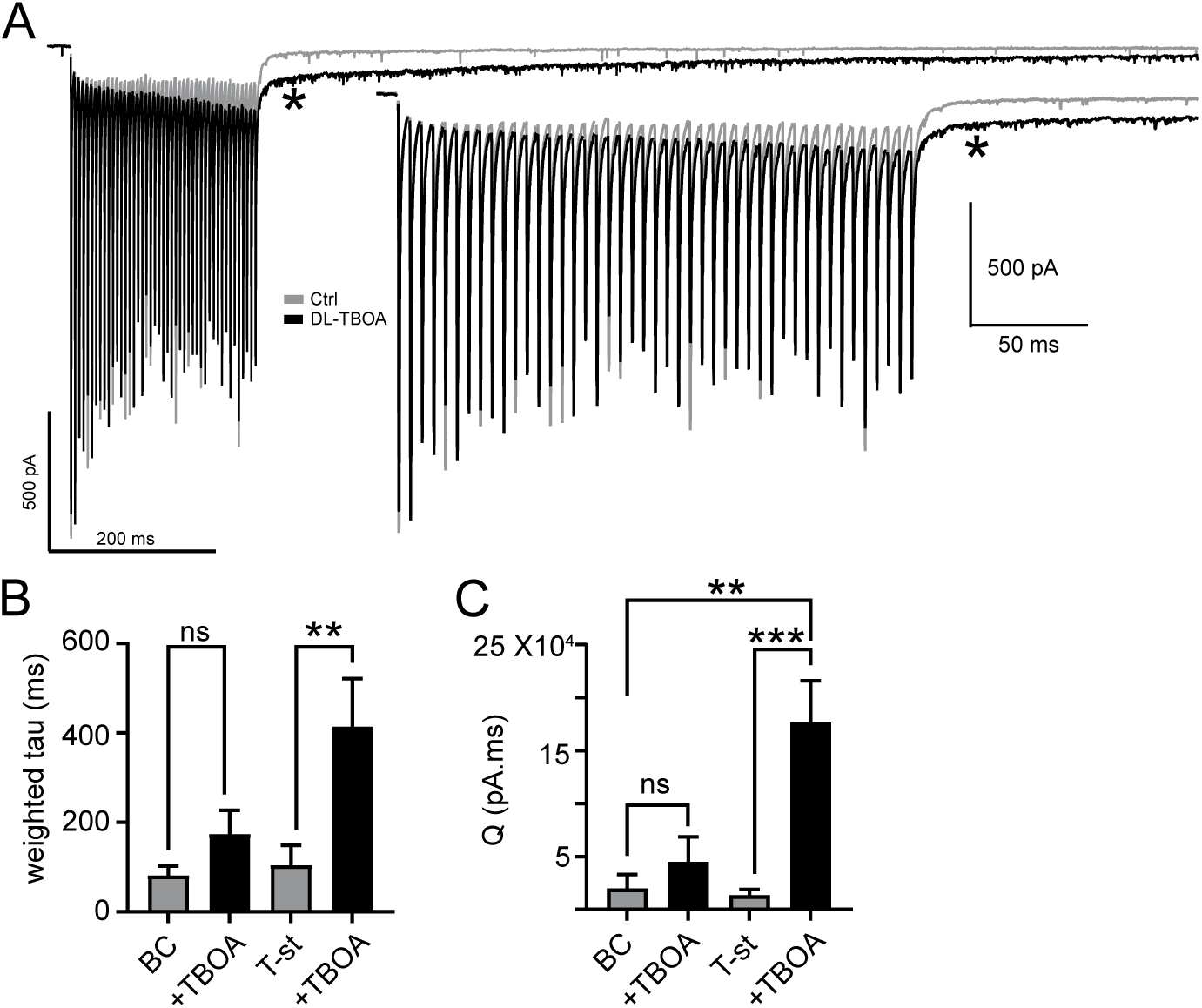
Effects of DL-TBOA on synaptic currents in bushy cells. **A.** Voltage-clamp traces of EPSC trains in a bushy cell, with and without DL-TBOA. Asterisk highlights longer decay of current in DL-TBOA. **B.** Comparison of weighted decay tau after EPSC train for bushy cells (BC) and T-stellate cells (T-st) with and without DL-TBOA. Increase in decay time in bushy cell with DL-TBOA did not reach significance. **C.** As in B, but for total charge during current decay phase ((*p* ** < 0.005, *** < 0.0005).

## Discussion

Plasma membrane glutamate transporters are considered essential for maintaining background extracellular glutamate at non-excitotoxic levels. Accordingly, complete block of transporter activity leads to dramatic buildup of glutamate (Sarantis et al., 1993; Yang and Xu-Friedman, 2009) and, as observed in this study, eventual loss of neural signaling. Additionally, the binding capacity of transporters can rapidly determine peak glutamate levels during synaptic transmission (Diamond and Jahr, 1997). However, it has not been clear what role transporters play in the ongoing dynamics of synaptic transmission and in the information coding dependent on that dynamic activity. Here, we titrated the magnitude of blockade so that background glutamate, assessed by membrane potential or holding current, remained low, and then examined the efficacy of synaptic transmission during stimuli in the physiological range. We found that while partial transporter block had little effect on single EPSCs, within only 2-3 presynaptic stimuli, glutamate accumulation occurred to such a degree that the temporal relationship between pre- and postsynaptic activity was severely compromised. Postsynaptic responses to brief high-frequency trains of presynaptic spikes, similar to normal activity of AN (Taberner and Liberman, 2005), led to prolonged postsynaptic firing when transporters were blocked, thereby distorting normal input-output relationships. Thus, this study demonstrates that a key feature of information coding in the auditory system, the tight relationship between pre-and postsynaptic signaling across time, is dependent not only on the rapid release of glutamate and its binding to AMPARs, but also to its rapid removal by EAATs.

Individual EPSCs exhibited an initial rapid current decay of less than a millisecond, likely due to rapid diffusion from the postsynaptic membrane, and this decay was unaffected by the presence of TBOA, indicating a prominent role for passive diffusion of transmitter. However, later phases of decay in the milliseconds time scale were slowed in TBOA, indicating that passive diffusion was unable to fully clear synaptic spaces of transmitter. This slowing of clearance led to buildup of transmitter near postsynaptic receptors that increased steadily with successive presynaptic spikes. That the slow current buildup reflected glutamate accumulation was confirmed by its sensitivity to AMPAR blockade: given the low affinity of AMPAR (Patneau and Mayer, 1990), accumulated glutamate must have continuously rebound and reactivated receptors for hundreds of milliseconds following cessation of exocytosis. Pharmacological experiments indicated that this accumulation of transmitter is prevented by the rapid action of both glial (EAAT1,2) and neuronal (EAAT3) transporters, consistent with the known expression of these transporters in cochlear nuclear astrocytes, T-stellate cells, as well as in spiral ganglion cells, which give rise to AN synapses (Jing et al., 2025; https://screen.hms.harvard.edu/harvard/). Where are these glial and neuronal transporters located with respect to AN synapses? AN fibers form bouton terminals on the somata, initial segment and dendritic shafts of T-stellate cells (Cant, 1981; Smith and Rhode, 1989b; Josephson and Morest, 1998), with glial cells covering and separating adjacent boutons. Given that 1) glial and neuronal transporters remove synaptic glutamate at AN-to-T-stellate cell synapses, 2) glutamate accumulation occurs after only a few spikes, and 3) that glutamate released from synapses made by different axonal inputs appears not to overlap, it is likely that transporters are distributed in glial cells overlying individual boutons and on pre- and postsynaptic structures as well. Such an arrangement would optimally isolate synapses and maintain glutamate at low levels between release events at individual synapses.

Little is known about how transporters contribute to information processing in the brain. Background glutamate levels are generally kept quite low by transporters (Herman et al., 2011; Chiu and Jahr, 2017), and it is assumed that such background transmitter would have little adaptive value in neural function. One striking exception is the large synaptic cleft formed by cerebellar mossy fibers and unipolar brush cell dendrites, where significant ambient glutamate persists despite normal transporter function. The tonic AMPAR and metabotropic glutamate receptor currents generated by this ambient glutamate enable the brush cells to mediate complex transformations of mossy fiber spike patterns that characterize vestibular signaling (Balmer et al., 2021). Transporter block distorted these transformations by enhancing background glutamate, prolonging the response to brief synaptic signals by over tenfold (Balmer et al., 2021). In the present study, the effects of transporters on ambient glutamate and stimulus evoked glutamate transients were separated by adjusting the concentration of TBOA, revealing that glutamate removal during brief rapid synaptic signaling is essential for the coding function in the auditory system.

Another cell-type-specific difference in transporter role was observed in the effects of TBOA on T-stellate cells and bushy cells, with T-stellate cells being far more compromised by transporter block. This difference was remarkable given that both cell types receive the same AN input, and that bushy cells receive particularly giant AN terminals that generate enormous synaptic currents (Yang and Xu-Friedman, 2009), and thus might be expected to have greater challenges in glutamate clearance. Yet bushy cell spike probability was little affected by transporter block, even when glutamate accumulation led to a steady depolarization of resting membrane potential (Yang and Xu-Friedman, 2015). Two key factors may contribute to this effect in bushy cells. First, the spherical shape of bushy cells and the concentration of synapses on the somatic rather than dendritic surface leads to greater non-neuronal space between adjacent bushy cells and enable more complete dilution of transmitter near the synapse (Spirou et al., 2023). Second, the presence of large, low-threshold K^+^ currents in bushy cells resist steady depolarizing currents and enable the neurons to maintain excitability despite glutamate accumulation (Cao et al., 2007).

By contrast, T-stellate cells have bouton synapses contributed by multiple AN fibers distributed along their dendrites within a dense neuropil (Cant, 1981; Smith and Rhode, 1989b; Josephson and Morest, 1998). Moreover, AN inputs to the cochlear nucleus are tonotopically organized, such that synapses within a narrow region are likely to be nearly synchronously active as they carry signals from narrow ranges of sound frequency. An added contributing feature may be the “synaptic nests” of cochlear nucleus which feature dense synaptic clusters, with EAATs distributed only at their edges (Josephson and Morest, 2003), although it is not certain if T-stellate cells contribute in such nests. Importantly, the voltage-sensitive conductances of T-stellate cells ensure summation, rather than shunting, of depolarizing currents. As a result, glutamate clearance by diffusion alone is apparently insufficient to prevent a massive buildup of transmitter near synapses resulting in protracted excitation as glutamate levels gradually fall. Transporter action at the synapse sites on T-stellate cells is therefore essential to allow individual AN inputs to contribute to postsynaptic responses in a linear fashion, and to enable the neurons to respond precisely and rapidly to changes in the number of active inputs and their frequency of firing.

Beyond their role in neural signaling, disruption of glutamate transporter function is a well-established contributing factor to many neurodegenerative diseases like amyotrophic lateral sclerosis (ALS), Alzheimer’s disease (AD), and Parkinson’s disease (Van Den Bosch et al., 2006; Arnold et al., 2024). For instance, in ALS, disrupted glutamate re-uptake, primarily due to dysfunctional EAAT2, leads to excessive glutamate accumulation and subsequent excitotoxic death of motor neurons. Glutamate excitotoxicity is also increasingly implicated in various auditory disorders. One example is that in noise-induced hearing loss, excessive glutamate accumulation can lead to loss of both inner and outer hair cells in the cochlea (Hakuba et al., 2000; Moser and Starr, 2016). Disruption of central glutamate signaling due to glutamate accumulation may contribute to hyperactivity and tinnitus. Thus, by showing that even a partial impairment of glutamate transport function can rapidly compromise information coding in the auditory system, our findings shed light on the importance of glutamate transporter function in normal neural signaling and potentially in known pathologies.

## Materials and Methods

### Key resources table

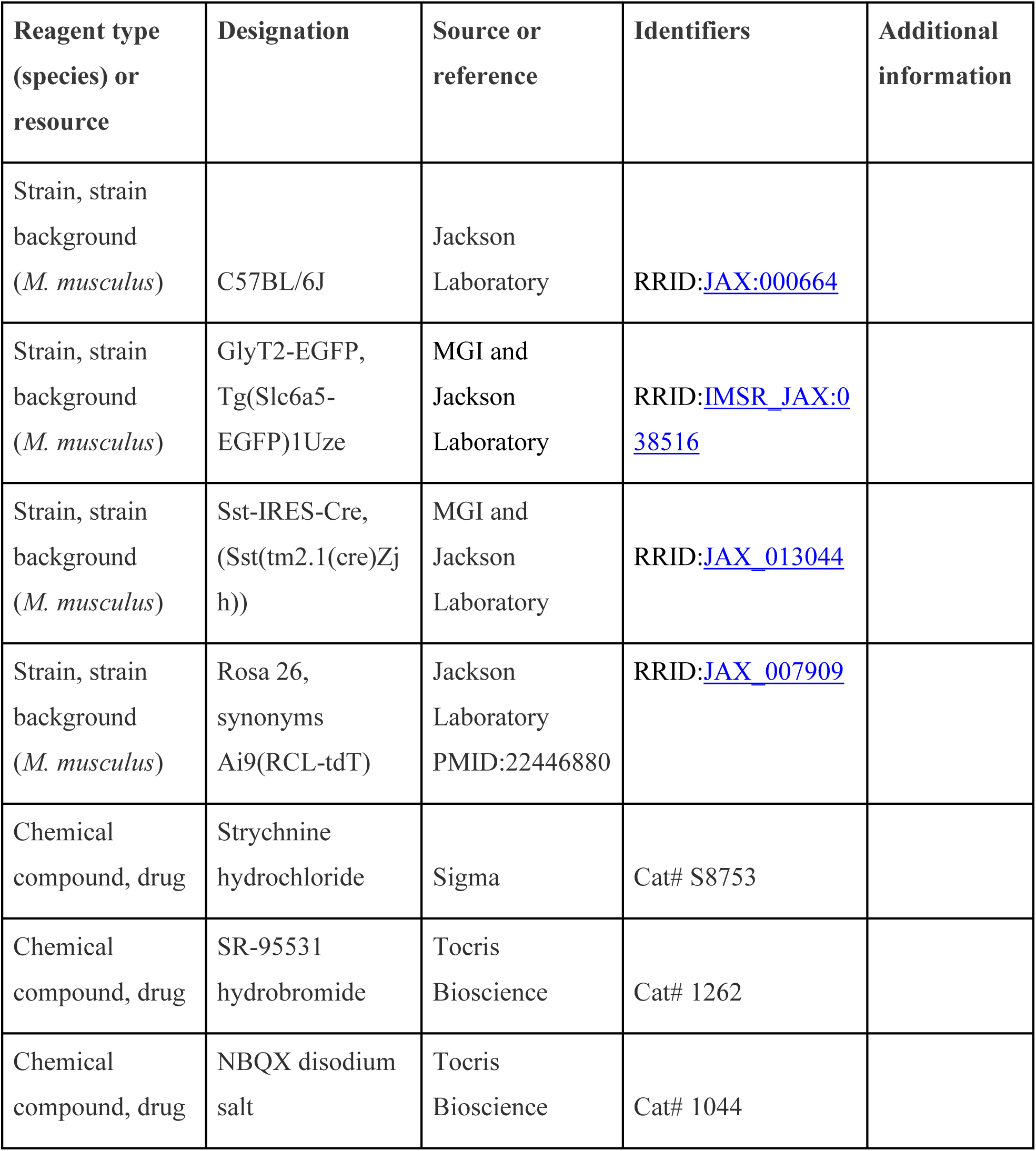
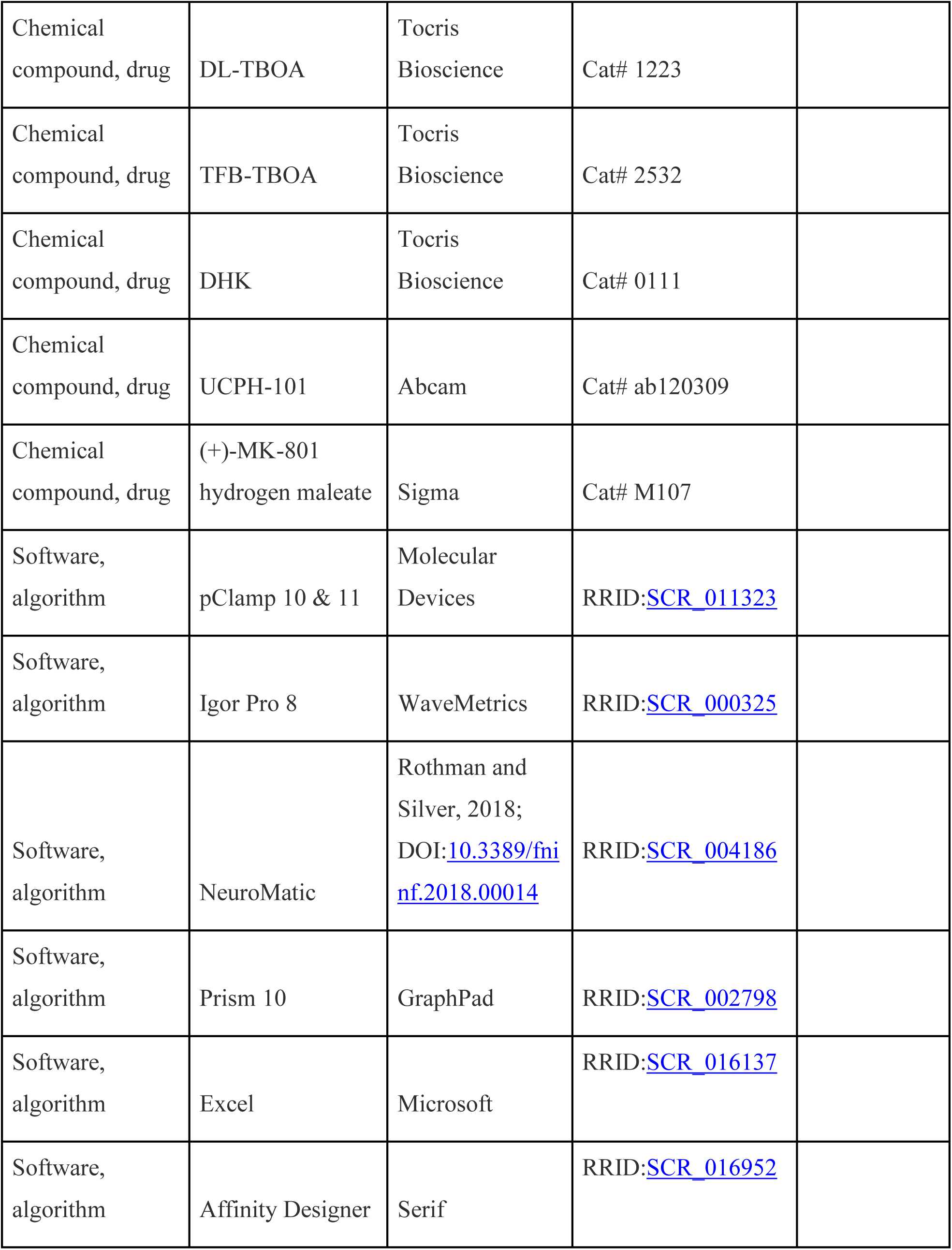

| Reagent type<br>(species) or<br>resource | Designation | Source or<br>reference | Identifiers | Additional<br>information |
| --- | --- | --- | --- | --- |
| Strain, strain<br>background<br>(*M. musculus*) | C57BL/6J | Jackson<br>Laboratory | RRID:JAX:000664 |  |
| Strain, strain<br>background<br>(*M. musculus*) | GlyT2-EGFP,<br>Tg(Slc6a5-EGFP)1Uze | MGI and<br>Jackson<br>Laboratory | RRID:IMSR_JAX:038516 |  |
| Strain, strain<br>background<br>(*M. musculus*) | Sst-IRES-Cre,<br>(Sst(tm2.1(cre)Zjh)) | MGI and<br>Jackson<br>Laboratory | RRID:JAX_013044 |  |
| Strain, strain<br>background<br>(*M. musculus*) | Rosa 26,<br>synonyms<br>Ai9(RCL-tdT) | Jackson<br>Laboratory<br>PMID:22446880 | RRID:JAX_007909 |  |
| Chemical<br>compound, drug | Strychnine<br>hydrochloride | Sigma | Cat# S8753 |  |
| Chemical<br>compound, drug | SR-95531<br>hydrobromide | Tocris<br>Bioscience | Cat# 1262 |  |
| Chemical<br>compound, drug | NBQX disodium<br>salt | Tocris<br>Bioscience | Cat# 1044 |  |
| Chemical compound, drug | DL-TBOA | Tocris Bioscience | Cat# 1223 |  |
| Chemical compound, drug | TFB-TBOA | Tocris Bioscience | Cat# 2532 |  |
| Chemical compound, drug | DHK | Tocris Bioscience | Cat# 0111 |  |
| Chemical compound, drug | UCPH-101 | Abcam | Cat# ab120309 |  |
| Chemical compound, drug | (+)-MK-801 hydrogen maleate | Sigma | Cat# M107 |  |
| Software, algorithm | pClamp 10 & 11 | Molecular Devices | RRID:SCR_011323 |  |
| Software, algorithm | Igor Pro 8 | WaveMetrics | RRID:SCR_000325 |  |
| Software, algorithm | NeuroMatic | Rothman and Silver, 2018;<br>DOI:10.3389/fninf.2018.00014 | RRID:SCR_004186 |  |
| Software, algorithm | Prism 10 | GraphPad | RRID:SCR_002798 |  |
| Software, algorithm | Excel | Microsoft | RRID:SCR_016137 |  |
| Software, algorithm | Affinity Designer | Serif | RRID:SCR_016952 |  |

### Animals

In this study, we used wild-type (C57BL/6J), GlyT2-EGFP (Zeilhofer et al., 2005), and somatostatin (Sst)-Cre tdTomato mice of either sex, from postnatal days 17–40 (P17–40). GlyT2-EGFP mice were backcrossed into the C57BL/6J line and maintained as heterozygous. Sst-tdTomato mice were generated by crossing homozygous Sst-IRES-Cre knock-in mice with homozygous Ai9 (RCL-tdTomato) reporter mice (Jackson Laboratory), enabling Sst-IRES-Cre-dependent expression of tdTomato. Mice were housed in the animal facility managed by the Department of Comparative Medicine at the Oregon Health and Science University. All procedures were approved by the Oregon Health and Science University’s Institutional Animal Care and Use Committee.

### Brain slice preparation

Animals were anesthetized with isoflurane and decapitated. The brain was quickly dissected out and immersed in ice-cold sucrose cutting solution containing the following (in mM): 76 NaCl, 26 NaHCO_3_, 75 sucrose, 1.25 NaH_2_PO_4_, 2.5 KCl, 25 glucose, 7 MgCl_2_, and 0.5 CaCl_2_, bubbled with 95% O2:5% CO2 (pH 7.8, 305 mOsm). Parasagittal cochlear nucleus sections, 180–210 μm in thickness were cut with a vibratome (7000smz-2, Campden Instruments) in ice-cold sucrose solution. Slices were transferred into standard artificial cerebrospinal fluid (ACSF) containing (in mM): 125 NaCl, 20 NaHCO_3_, 1.2 KH_2_PO_4_, 2.1 KCl, 3 Na-HEPES, 10 glucose, 1 MgCl_2_, 1.2 CaCl_2_, 2 Na-pyruvate, and 0.4 Na-L-ascorbate, bubbled with 95% O2:5% CO2 (pH 7.4, 300–310 mOsm). Slices recovered at 34°C for 40 min and were maintained at room temperature until recording.

### Electrophysiological recordings

Slices were transferred to a recording chamber and perfused with standard ACSF using a peristaltic pump (Ismatec) at 2–3 ml/min and maintained at ∼34°C with an inline heater (TC0324B, Warner Instruments). Cells were viewed using an Olympus BX51WI microscope with a 60X objective, equipped with an infrared Dodt contrast, a CCD camera (IR- 2000, DAGE-MTI), and fluorescence optics. In slices from GlyT2-GFP mice, GFP-negative cells in the VCN were targeted for T-stellate and bushy cells whereas in slices from Sst-TdTomato mice, tdTomato-positive neurons were targeted from VCN for T-stellate and bushy cells recordings. Sutter Lambda FLEDs with 480 nm and 530 nm high-power LEDs were used to visualize fluorescent label cells. Recording pipettes were pulled from 1.5 mm OD, 0.84 mm ID borosilicate glass (WPI-1B150F-4) to a resistance of 2–4 MΩ using a horizontal puller (Sutter Instrument P1000).

The internal recording solution contained in (mM): 113 K gluconate, 2.75 MgCl_2_, 1.75 MgSO_4_, 0.1 EGTA, 14 Tris-phosphocreatine, 4 Na_2_-ATP, 0.3 Tris-GTP, 9 HEPES with pH adjusted to 7.25 with KOH, mOsm adjusted to 290 with sucrose (E_Cl_-84 mV). Whole-cell patch-clamp recordings were made using a Multiclamp 700B amplifier and pClamp 10.7 or 11.3 software (Molecular Devices). Voltages were corrected for a junction potential of −10 mV. Signals were digitized at 20–40 kHz and filtered at 10 kHz by Digidata 1440A (Molecular Devices). In voltage-clamp mode, cells were held at −60 to −65 mV, with access resistance of 5‒20 MΩ compensated to 50–60%. In current-clamp mode, the resting membrane potential was maintained between −60 to −70 mV with bias current. All experiments were conducted in the presence of inhibitory synaptic blockers. For AN electrical stimulation, a concentric bipolar electrode (CBBPC75, FHC) was placed in the AN root. Electric shock pulses (10–250 μs duration, 0–60 V) were delivered using a stimulus isolation unit (Iso-Flex, A.M.P.I).

### Analysis

The decay of the averaged EPSC current was analyzed in Clampfit (10.7 and 11.2) and fitted with a multiple exponential components using the simplex algorithm. Adequacy of fit was judged by eye, or in less obvious cases, by the criterion of halving of sum-squared error when an additional exponential component was added. When 4-5 exponential components were needed, the slowest component was fitted separately, and that exponential value was fixed (unchanged) during the multiexponential fitting procedure. Weighted decay tau was calculated by scaling each exponential time constant by the fractional contribution of its exponential term to the overall fit.

### Electrophysiological and statistical analyses

Electrophysiological data were analyzed using pClamp (10.4 or 11.2, Molecular Devices), IGOR Pro v8 (WaveMetrics), and NeuroMatic (Rothman and Silver, 2018). Figures were made using IGOR Pro, and Affinity Designer. Statistics were performed in IGOR Pro, Microsoft Excel, or Prism10 (GraphPad). For statistical analysis, groups were tested for normality using Shapiro-Wilk test before using parametric test. One-way or two-way ANOVA with repeated measures were used for comparisons across different groups followed by post hoc comparison test. Paired and unpaired *t*-tests were used to compare means. The Friedman test or Kruskal-Wallis test with Dunn’s multiple comparison test were used for nonparametric distributions. Error bars are represented as mean ± SEM unless otherwise stated.

Dataset can be found at OSF: https://osf.io/58h7c

## Notes

### Competing Interest Statement

The authors have declared no competing interest.

